## Supplementary material for "Comprehensive analysis of male-free reproduction in *Monomorium triviale* (Formicidae: Myrmicinae)": S1_File

**Materials and methods**

To confirm reproductive ability of *M. triviale* workers, we dissected a total of 100 intranidal (i.e., relatively young) workers (80 from 8 nests collected in locality no. 1 in Table 1 and 20 from 2 nests collected in locality no. 3). Each worker was first immobilized by soaking in 70% ethanol for 3 min. The body was then transferred to a 30-mm petri dish filled with distilled water, and the internal organs were pulled out from the end of the abdomen with precision forceps under a binocular microscope (SZ40; OLYMPUS Optical, Tokyo, Japan). Finally, we checked the absence of ovaries and the presence of other organs (e. g., crop, midgut, rectum and poison gland) in the worker abdomen.

**Results**

We successfully dissected 95 out of 100 individuals. The workers had no ovaries on the position homologous to the queens, indicating that they were obligatorily sterile (**Fig S1**).

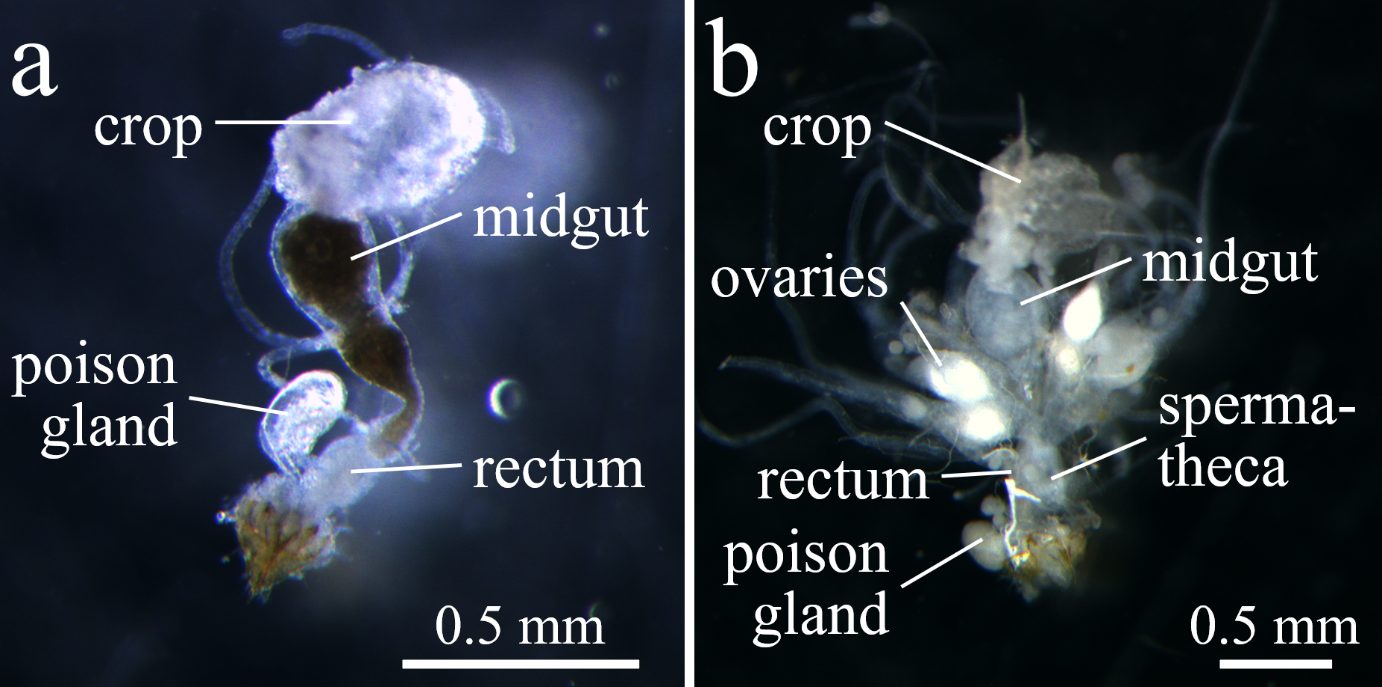

**Fig S1. Internal organs of (a) worker and (b) queen of *M. triviale***. Both workers and queens (individuals photographed were both from nest Mtri20200716_3) possessed normal digestive systems (crop, midgut and rectum) and poison glands, but only the queen had reproductive organs (ovaries and spermatheca).
